# PERSPECTIVES FOR POPULATION MANAGEMENT OF FELIDS IN INDIAN ZOOS

**DOI:** 10.1101/2022.09.18.508402

**Authors:** Lakshminarasimha R, Gowri Mallapur, Sonali Ghosh, Satya Prakash Yadav

## Abstract

The family: Felidae (hereafter referred to as felids) is among the commonly represented species in animal collections in Indian zoos. Of the globally recognised 45 species, 15 species (>30%) are housed in Indian zoos. Since 2007, the Central Zoo Authority has laid emphasis on ex situ conservation for seven threatened species by initiating planned breeding programmes.

We investigated the demographics of felids housed in Indian zoos using data from CZA annual inventory records. Between 1995-96 and 2019-20, the population of large felids have remained stable with a mean growth rate (λ) of 1.01; whereas the population of small felids have a marginally higher mean growth rate (λ) of 1.03. We further use Sustainability-index analysis to investigate whether the observed growth patterns arise from intrinsic (i.e. births/deaths) or extrinsic (acquisition/disposal) factors.

The management of felids in Indian zoos requires careful consideration of many factors including space, hybridisation, lack of pedigree knowledge, addition of wild-rescued specimens and colormorphs. We provide the first insights on how felids populations have fared at the family-level and species-level based on analysis of longitudinal data. The said analysis intends to inform plans to manage felid collections in Indian zoos. It should further present an outlook and also guide ongoing planned breeding programs of felids. Given the relatively large collection size and the corresponding conservation attention accorded to felids, our analysis will aid in setting priorities for collection planning, conservation education messaging, integration of in situ and ex situ efforts in the context of IUCN One Plan Approach.

## Introduction

The contribution of zoos to wildlife conservation has been emphasized in the recent years (IUCN SSC 2014). This includes providing animals for reintroductions (Gilbert et al. 2017), field-based monitoring and research (Che-Castaldo et al. 2018), and management of ex situ animal populations (Balmford et al. 1996; Conway 2011). An integrated approach is emphasised, that promotes an active contribution of lessons learnt from ex situ conservation to in situ conservation (Redford et al. 2012; IUCN SSC 2014). To be able to successfully realize the conservation goals, ex situ populations should be demographically and genetically stable (Lees and Wilcken 2009; Ballou et al. 2010). To this end, zoos have coordinated the breeding and exchange of animals among facilities through coordinated breeding programs since the 1980s (Ballou et al. 2010; Ballou and Traylor-Holzer 2011).

Central Zoo Authority (CZA) is an apex statutory body that recognizes and regulates operation of zoos in the country. Among other functions, the CZA coordinates breeding programs for threatened species and acquisition/transfer of animals between zoos, thereby, regulating animal collections in zoos. The overall goal is to ensure that collections are demographically robust and genetically representative of wild counterparts for the foreseeable future.

Standardised inventory information maintained by zoos offers insights on dynamics of zoo populations. Recognised zoos submit inventory of collections detailing acquisitions, births, disposals and deaths annually to the Central Zoo Authority.

In this paper, we analyse longitudinal inventory data of species belonging to the family *Felidae* housed in Indian zoos between 1995 and 2020. The analysis is an attempt to get an overview of population trends, growth rates and holding patterns of felids. The results of this retrospective analysis whould provide perspectives for managing felid populations in Indian zoos sustainably.

## Methods

In this study, we selected all the species belonging to the family *Felidae* housed in Indian zoos (Table 1). The data pertaining to these species was obtained from inventory records submitted by recognised zoos to the Central Zoo Authority, subsequently digitised and publicly available (www.cza.nic.in). The inventory was available for each fiscal year i.e., from 1^st^ April of a given year to the 31^st^ March of the following year. It comprised of species-wise opening and closing stock including details of births, acquisitions, disposals and deaths for each fiscal year. In total, data from 143 zoos housing one or more target species for at least one fiscal year was used for the study.

**Table 1.** Species and data availability used in the current study.

| No. | Species | Data availability on housing | Category |
| --- | --- | --- | --- |
| 1 | African Lion *Panthera leo* | Apr 1, 1995 – Mar 31, 1996 to Apr 1, 2019 – Mar 31, 2020 | Large felids |
| 2 | Asiatic Lion *Panthera leo persica* | Apr 1, 1995 – Mar 31, 1996 to Apr 1, 2019 – Mar 31, 2020 |  |
| 3 | Bengal Tiger *Panthera tigris* | Apr 1, 1995 – Mar 31, 1996 to Apr 1, 2019 – Mar 31, 2020 |  |
| 4 | Jaguar *Panthera onca* | Apr 1, 1995 – Mar 31, 1996 to Apr 1, 2019 – Mar 31, 2020 |  |
| 5 | Leopard *Panthera pardus* | Apr 1, 1995 – Mar 31, 1996 to Apr 1, 2019 – Mar 31, 2020 |  |
| 6 | Puma *Puma concolor* | Apr 1, 1995 – Mar 31, 1996 to Apr 1, 2009 – Mar 31, 2010 |  |
| 7 | Siberian Tiger *Panthera tigris altaica* | Apr 1, 1995 – Mar 31, 1996 to Apr 1, 2013 – Mar 31, 2014 |  |
| 8 | Snow Leopard *Panthera uncia* | Apr 1, 1995 – Mar 31, 1996 to Apr 1, 2019 – Mar 31, 2020 |  |
| 9 | South African Cheetah *Acinonyx jubatus* | Apr 1, 2008 – Mar 31, 2009 to Apr 1, 2019 – Mar 31, 2020 |  |
| 10 | Sumatran Tiger *Panthera tigris sumatrae* | Apr 1, 2016 – Mar 31, 2017 to Apr 1, 2018 – Mar 31, 2019 |  |
| 11 | Clouded Leopard *Neofelis nebulosa* | Apr 1, 1995 – Mar 31, 1996 to Apr 1, 2019 – Mar 31, 2020 | Small felids |
| 12 | Asian Golden Cat *Catopuma temminckii* | Apr 1, 1995 – Mar 31, 1996 to Apr 1, 2019 – Mar 31, 2020 |  |
| 13 | Serval *Leptailurus serval* | Apr 1, 2017 – Mar 31, 2018 to Apr 1, 2019 – Mar 31, 2020 |  |
| 14 | Caracal *Caracal caracal* | Apr 1, 2016 – Mar 31, 2017 to Apr 1, 2019 – Mar 31, 2020 |  |
| 15 | Leopard Cat *Prionailurus bengalensis* | Apr 1, 1995 – Mar 31, 1996 to Apr 1, 2019 – Mar 31, 2020 |  |
| 16 | Rusty-spotted Cat *Prionailurus rubiginosus* | Apr 1, 1995 – Mar 31, 1996 to Apr 1, 2019 – Mar 31, 2020 |  |
| 17 | Fishing Cat *Prionailurus viverrinus* | Apr 1, 1995 – Mar 31, 1996 to Apr 1, 2019 – Mar 31, 2020 |  |
| 18 | Jungle Cat *Felis chaus* | Apr 1, 1995 – Mar 31, 1996 to Apr 1, 2019 – Mar 31, 2020 |  |
| 19 | Hybrid Lion *Panthera leo* | Apr 1, 1995 – Mar 31, 1996 to Apr 1, 2019 – Mar 31, 2020 | Hybrid and colour morph specimens |
| 20 | Leucistic Tiger *Panthera tigris* | Apr 1, 1995 – Mar 31, 1996 to Apr 1, 2019 – Mar 31, 2020 |  |
| 21 | Melanistic Leopard *Panthera pardus* | Apr 1, 1995 – Mar 31, 1996 to Apr 1, 2019 – Mar 31, 2020 |  |

The nature of the data available i.e. basic inventory with records of births, deaths, acquisitions and disposals was apt for conducting S-index analysis (Lynch 2018). We computed the following S-index metrics for species mentioned in Table 1, where N = number of individuals in the collection, t = time step, B = births, D = deaths, I = imports (acquisitions) to the collection, and E = exports (dispositions) from the collection.

1. Total lambda (λ_T_) = The proportional change in population size from one year to the next, calculated as:

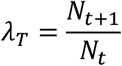

2. Intrinsic lambda (λ_I_) = The proportional change in population size due to Births/Hatches and Deaths, calculated as:

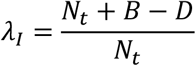

3. Extrinsic lambda (λ_E_) = The proportional change in population size due to Acquisitions and Disposals, calculated as:

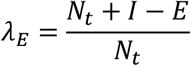

Since Sumatran Tiger *Panthera tigris ssp. sondaica*, Cougar *Puma concolor*, Siberian Tiger *Panthera tigris ssp. altaica*, Serval *Leptailurus serval*, Caracal *Caracal caracal* and Asian Golden Cat *Catopuma temminckii*, Leopard (melanistic) *Panthera pardus* are either not housed in Indian zoos anymore or data includes fewer than 10 individuals, only descriptive account of the population trends is presented. For the remaining species, the S-index metrics were computed for each fiscal year and summarised for investigating patterns of population growth.

All the analyses were performed using R Statistical Program (R Core Team 2020) using the packages ‘tidyverse’ (Wickham et al., 2019), ‘ggpubr’ (Kassambara, 2020) and ‘viridis’ (Garnier et al., 2021).

## Results

### (a) Population trends and housing patterns

In total data from 143 Indian zoos housing felids between 1^st^ April 1995 to 31^st^ March 2020 including 14852 observations of births, acquisitions, disposals and deaths were analysed. The species-wise census trends of felids species in captivity and corresponding number of holding institutions is indicated in Figure 1.

**Figure 1.**
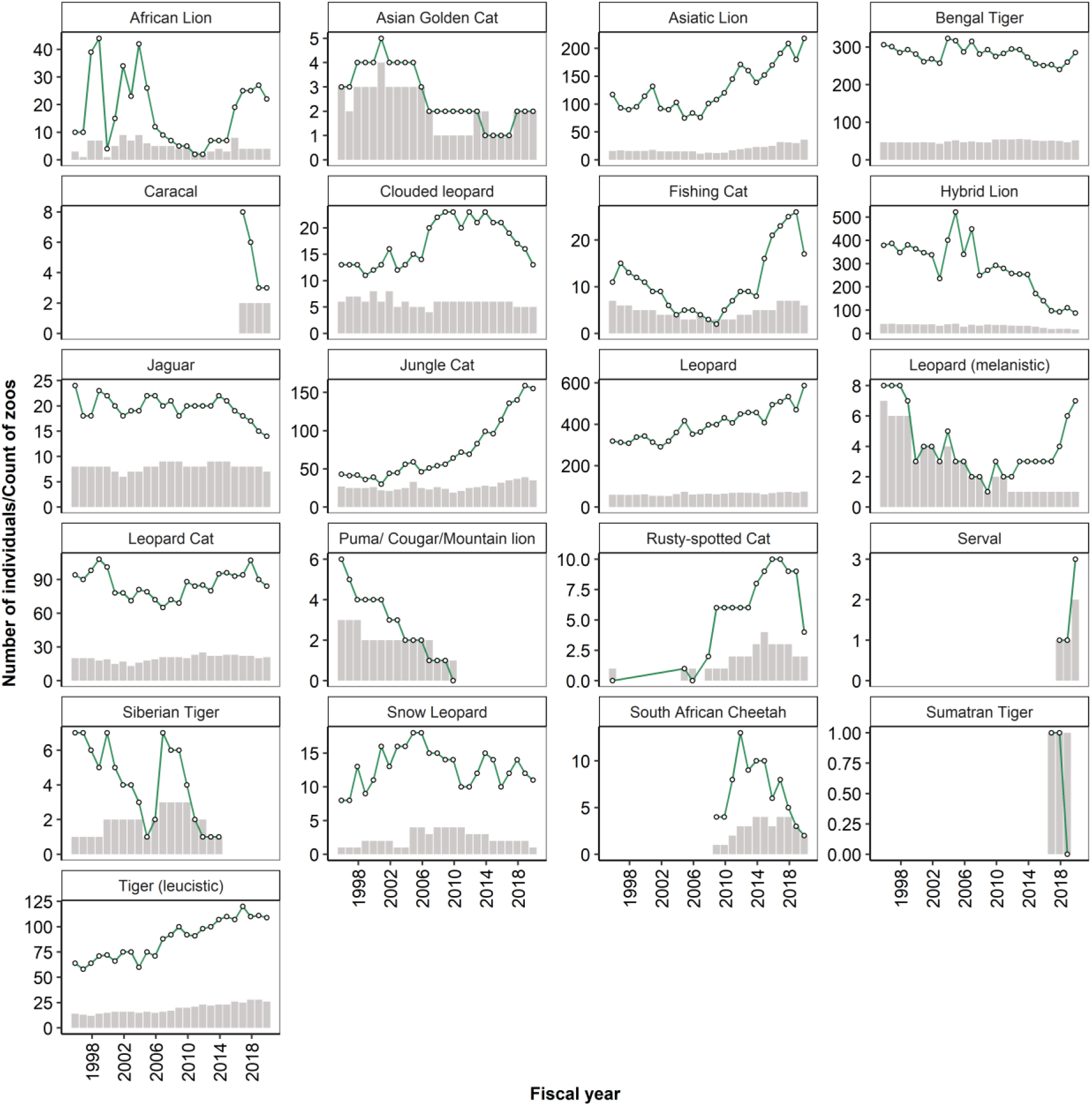
Census of felids housed in Indian zoos: 1995-96 to 2019-20. Census details are collated for fiscal years i.e. from 1^st^ April of a given year to the 31^st^ March of the following year. The line indicates the total number of males, females and unsexed individuals housed in all the zoos at the end of each fiscal year. Bars represent the number of zoos housing the species in the respective fiscal year.

The captive population of *viz*. Leopard *P*.*pardus*, Asiatic Lion *P*.*leo ssp. persica* and Jungle Cat *F. chaus* has steadily increased and has more than doubled during the said time period. The captive population of Bengal Tiger *P*.*tigris ssp. tigris*, Leopard Cat *P*.*bengalensis* has remained relatively stable. Non-native species *viz*. Puma *P. concolor*, Sumatran Tiger *P. tigris ssp. sumatrae* and Siberian Tiger *P. tigris altaica* were housed briefly and are no longer housed in any zoos. All the remaining felid species have continued to remain as small collections without significant growth. The captive population of Bengal Tiger *P*.*tigris* (leucistic) has steadily increased, whereas Hybrid lion *P. leo ssp*. has progressively decreased. Leopard *P. pardus* (melanistic) has continued to remain as a small collection without any significant growth.

The holding patterns by the zoos have remained consistent over the years across species (Figure 1). As on 31^st^ March, 2021, 85 zoos are housing at least one of the species listed in Table 1. Large felids *viz*. Asiatic Lion *P. leo ssp. persica*, Bengal Tiger *P*.*tigris ssp. tigris*, Leopard *P. pardus* and small felids Jungle Cat *F. chaus* and Leopard Cat *P*.*bengalensis* are the most commonly housed species in Indian zoos. All the remaining felid species are housed as small collections in few zoos in comparison to hybrid and color morph specimens that have a relatively higher representation. The distribution of felids across zoos at the end of fiscal year 2019-2020 is represented in Figure 2.

**Figure 2.**
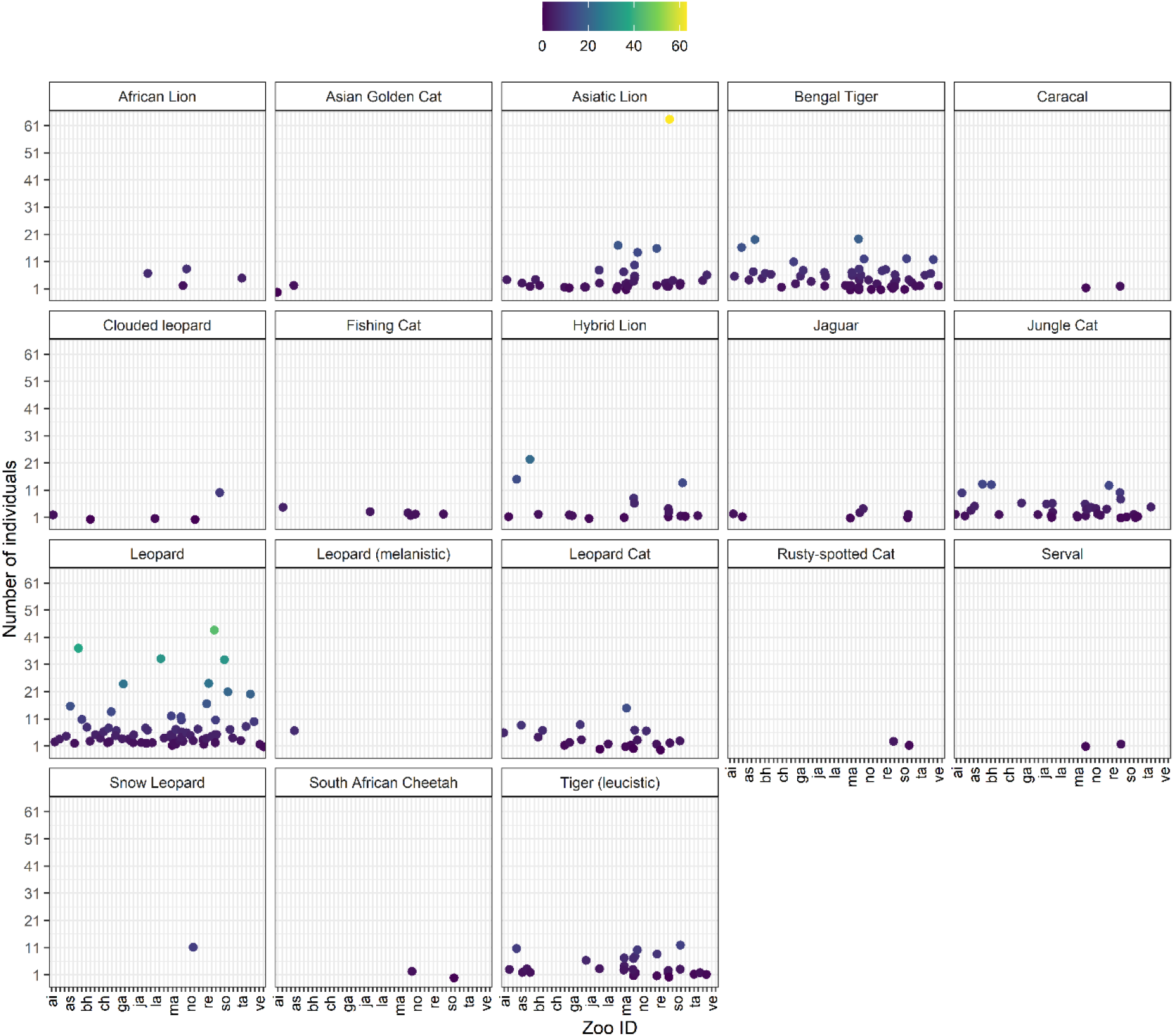
Housing patterns of felids in Indian zoos at the end of 2019-20 fiscal year. Zoo ID’s are abbreviations derived from zoo names. Colour gradient represents the number of individuals in respective zoo.

For each fiscal year, the proportion of acquisitions, births, disposals and deaths were computed and plotted across time to investigate their patterns (Figure 3). Acquisitions have been predominant in *P*.*pardus* and *F*.*chaus* and *P*.*bengalensis*; further, the birth:death ratio was less than zero in these species and also in the case of *N*.*nebulosa, P*.*uncia* and *P*.*t ssp. tigris*.

**Figure 3.**
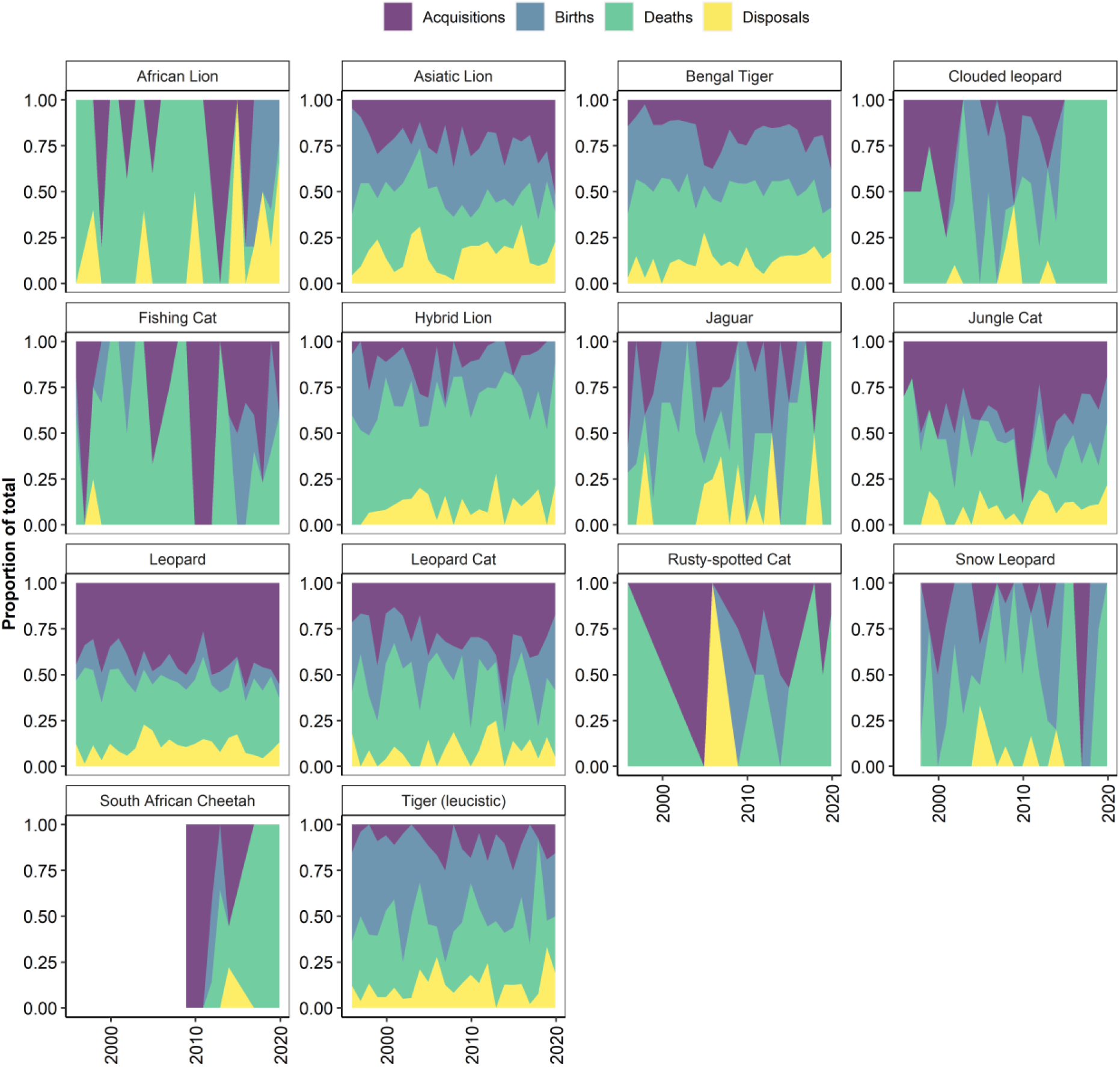
Proportion of acquisitions, births, deaths and disposals in select felids from 1995-96 to 2019-20.

We fitted a localised polynomial regression (loess) to investigate trends in collection sizes across years for select species of felids (Figure 4). The populations of *P. leo ssp. persica, P. viverrinus, P*.*pardus, F*.*chaus, P*.*tigris* (leucistic) show increasing trends, while the remaining species show variable, yet, declining trends.

**Figure 4.**
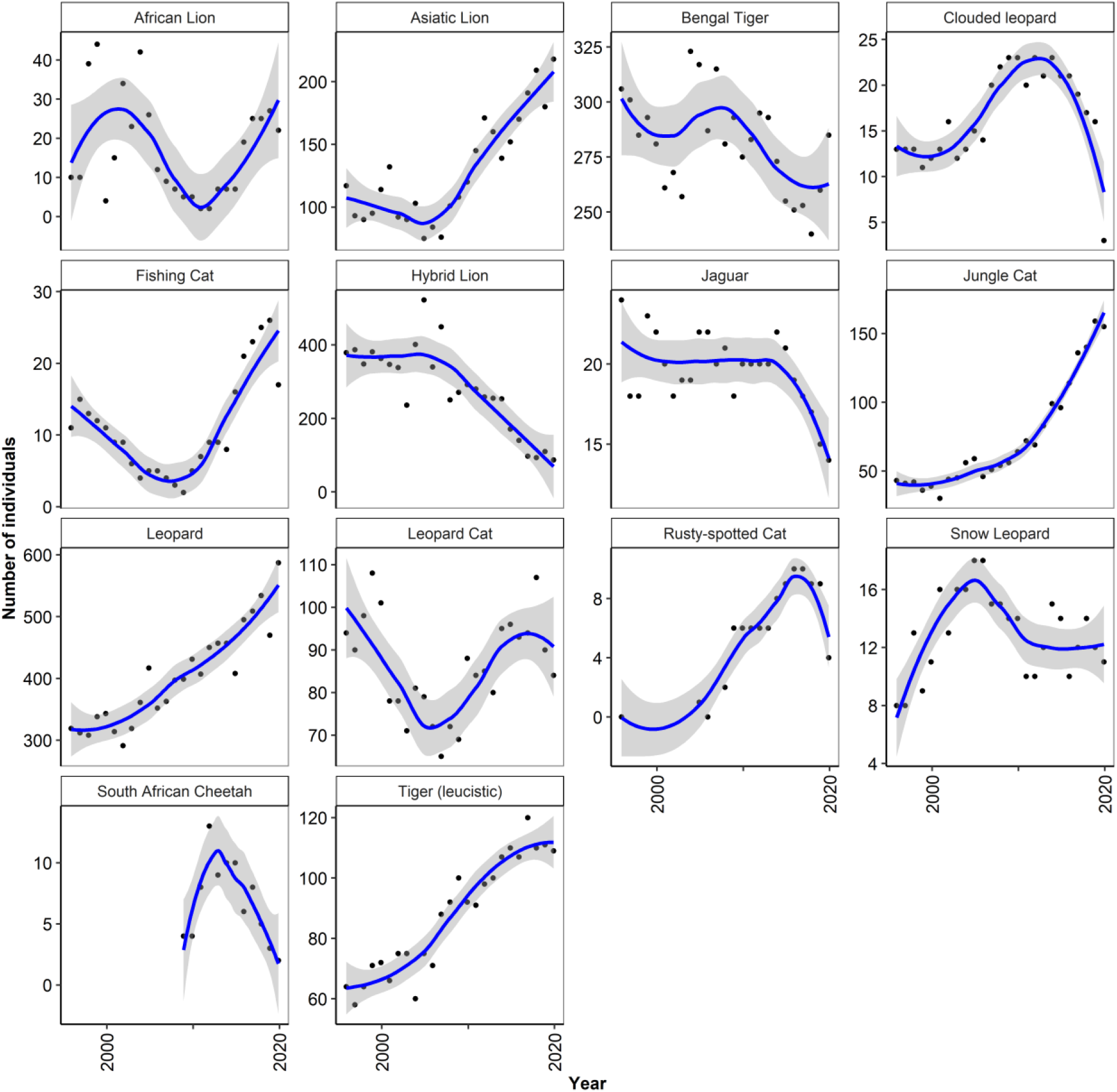
Trends (with loess smoothing and pointwise confidence intervals) of population sizes of select species of felids housed in Indian zoos from 1995-96 to 2019-20.

### (b) Total growth rate, intrinsic and extrinsic lambda

The mean growth rate across species is 1.05 (range = 0.97 – 1.36). To investigate changes in population growth across species, we calculated annual growth rates for each fiscal year and summarised it across species. Realised annual growth rates were highly variable across years (Figure 5). However, in general, growth rates remained close to replacement rate (growth rate [λ] ≥1). To understand drivers of the realised growth rates across species, we calculated intrinsic lambda (λ_I_) and extrinsic lambda (λ_E_) for each species. The values of λ_E_ was significantly higher than λ_I_ in all the species except Snow Leopard *P*.*uncia* and Bengal Tiger *P*.*tigris* (leucistic) (Figure 6). In the case of *P*.*unica* both the parameters had nearly equal values, whereas, in the case of *P*.*tigris* (leucistic), λ_I_ was significantly higher than λ_E_, indicating an intrinsically driven increase in the collection size.

**Figure 5.**
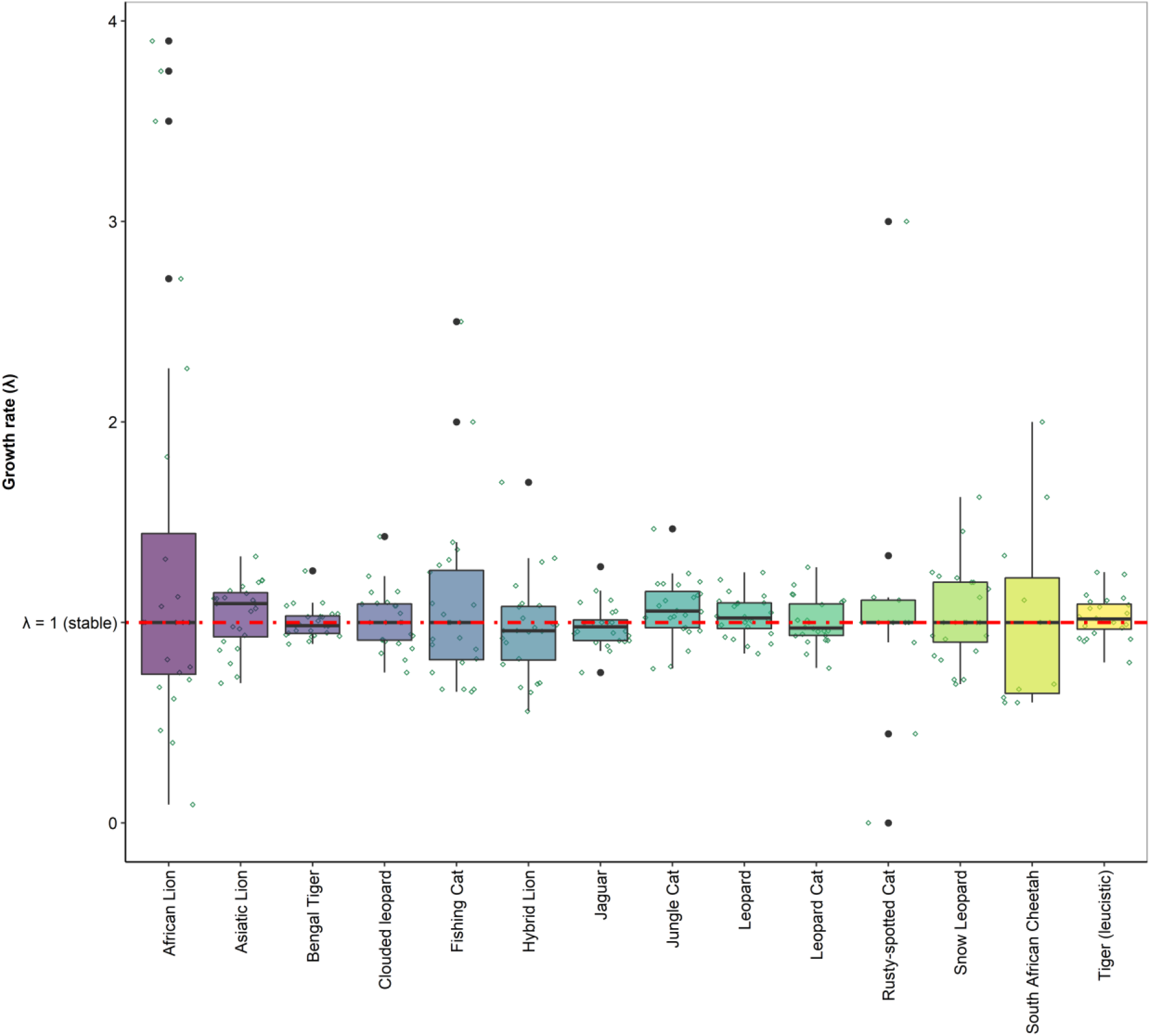
Species-wise growth rate (lambda [λ]) calculated from annual census data of felids housed in Indian zoos from 1995-96 to 2019-20. Red line indicates the stable growth rate. Data points in green indicate distribution of annual growth rates.

**Figure 6.**
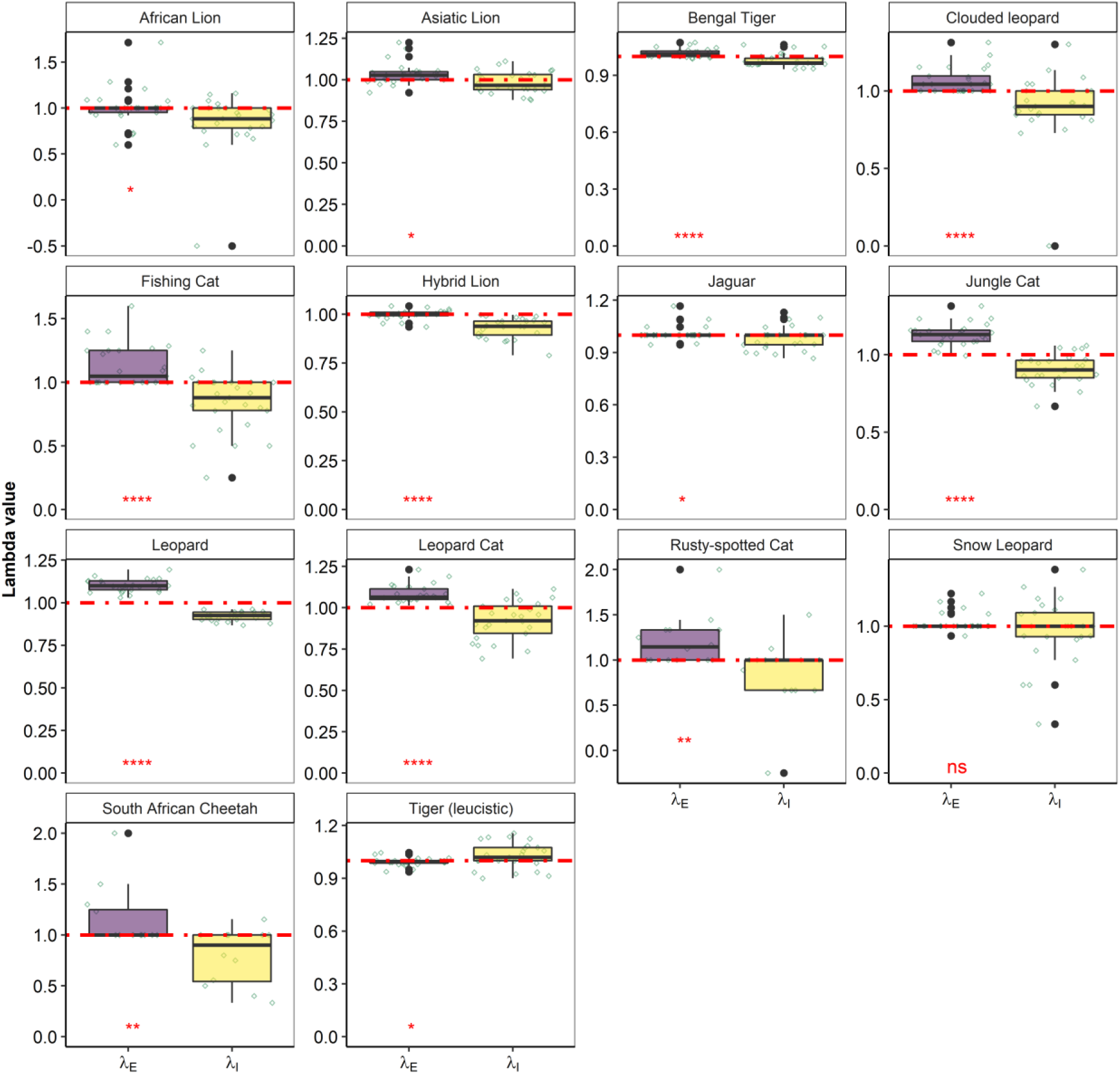
Species-wise intrinsic growth rate (λ_I_) and extrinsic growth rate (λ_E_) calculated from annual inventory records of felids housed in Indian zoos from 1995-96 to 2019-20. Red line indicates the stable growth rate. Data points in green indicate distribution of annual growth rates. Asterisk in red indicates significance levels, *ns* is non-significant.

## Discussion

Our analysis provides the first insights on long-term trends of felid populations in Indian zoos. While more than 50% of the zoos in the country continue to house one or more felids from this study, very few species demonstrate signs of long-term sustainability. Non-native species including *P*.*t ssp. sondaica, P. concolor, P*.*t ssp. altaica* are not housed in any zoos, whereas others such as *P*.*onca, L. serval, P. leo* and *A*.*jubatus* continue to remain as small collections without significant growth. Comparatively, native species have fared better. However, majorly, they too are housed as small collections in a few zoos. The trends of species that are housed in sizeable numbers in zoos are variable indicating fluctuating breeding successes correlated with a birth:death ratio of less than zero. Generally, for species housed in large numbers the growth rate is indicative of an increasing population. However, given the significantly higher rate of extrinsic lambda λ_E_ observed in most species, growth of the collection is therefore driven by acquisitions/disposals within the collection.

It is essential that demographic and genetic characteristics of collections are frequently evaluated to ensure that the population characteristics are aligned with the overall goal (e.g. insurance, display and education, conservation breeding etc.). Conversely, it is also important that goals are met, and outputs assessed, and results of the evaluation are used to adaptively manage populations.

The results presented in this paper offer long-term trends that can be adaptively used to manage collections and drive policy-level decisions addressing the following aspects:

1. Several species of felids are still housed as small collections with fewer than 20 individuals. Since, small populations are more susceptible to demographic stochasticity, genetic drift, and the effects of inbreeding depression (Lacy 1997; Ballou et al. 2010). The future management should achieve breeding success and promote population growth to shield them from these effects.
2. Several species (e.g. *P*.*tigris spp. tigris, P*.*rubiginosus*) are species prioritised for coordinated breeding by the Central Zoo Authority. The results of this analysis can be used to formulate conservation breeding plans for the identified species. Further, the results should also inform identification of felid species for initiation of conservation-oriented captive breeding programs.
3. This analysis provides a broad indication of species that can be successfully housed and bred (especially native species). It also presents how collections in zoos have varied over the years, thus allowing informed decisions pertaining to species acquisitions, movement of animals between zoos, management of rescued animals and in determining surplus collections.
4. The results presented here should serve as a baseline for further analysis and precise understanding of the factors that drive changes in collection size that will allow management of animal collections for sustainability.

## Acknowledgements

We like to thank zoo operators/directors for submission of Annual Inventory information of respective zoos. We would like to thank officials and staff of the CZA Secretariat, especially Vivek Goel, for curation and maintenance of inventory datasets, and Sruthy Boopathy for assistance is digitisation of inventory records.

